# DeepSeqPan, a novel deep convolutional neural network model for pan-specific class I HLA-peptide binding affinity prediction

**DOI:** 10.1101/299412

**Authors:** Zhonghao Liu, Yuxin Cui, Zheng Xiong, Alierza Nasiri, Ansi Zhang, Jianjun Hu

## Abstract

Interactions between human leukocyte antigens (HLAs) and peptides play a critical role in the human immune system. Accurate computational prediction of HLA-binding peptides can be used for peptide drug discovery. Currently, the best prediction algorithms are neural network based pan-specific models, which take advantage of the large amount of data across HLA alleles. However, current pan-specific models are all based on the pseudo sequence encoding for modeling the binding context and depend on the available HLA protein-peptide bound structures. In this work, we proposed a novel deep convolutional neural network model (DCNN) for HLA-peptide binding prediction, in which the encoding of the HLA sequence and the binding context are both learned by the network itself without requiring the HLA-peptide bound structure information. Our DCNN model is also characterized by its binding context extraction layer and dual outputs with both binding affinity output and binding probability outputs. Evaluation on public benchmark datasets shows that our DeepSeqPan model without HLA structural information in training achieves state-of-the-art performance on a large number of HLA alleles with good generalization capability. Since our model only needs raw sequences from the HLA-peptide binding pairs, it can be applied to binding predictions of HLAs without structure information and can also be applied to other protein binding problems such as protein-DNA and protein-RNA bindings. The implementation code and trained models are freely available at https://github.com/pcpLiu/DeepSeqPan.

## Introduction

Human leukocyte antigens (HLAs) are major histocompatibility complex (MHC) proteins located on the cell surface in human. HLAs play a critical role helping our immune system recognizing pathogens by binding to peptide fragments derived from pathogens and exposing them on the cell surface for recognition by appropriate T cells. Study of the binding mechanism between peptides and HLAs can help improve our understanding of human immune system and boost the development of protein-based vaccines and drugs [12,13]. Out of all classes of HLAs, we are interested in two major classes: class I and II. Class-I HLAs bind to peptides inside the cell while class-II HLAs bind to peptides outside the cell.

A big challenge of determining peptides binding to HLAs is the high polymorphism of HLA genes. As of March 2018, there are more than 17000^1^ HLA alleles deposited in the IMGT/HLA database. Experimentally testing the binding between peptides and different types of HLAs is costly and time-consuming. As a result, computational methods have been proposed to address this problem as more and more in vitro binding affinities data are published in databases such as IEDB [24], SYEPEITHI [18] and MHCBN [10].

Generally, current computational methods for peptide-HLA binding affinity prediction can be grouped into two categories: allele-specific and pan-specific models [4,6,8,11-15,17,25]. Allele-specific models are trained with only the binding peptides tested on a specific allele and a separate allele-specific binding affinity prediction model is needed for each HLA allele. NetMHC [12] and SMM [17] are the top allele-specific MHC binding prediction models. These models have the advantage of good performance when sufficient number of training peptide samples are available. However, due to the high polymorphism, for many HLA alleles, there are no or just a few experimentally determined binding affinity data. To address this data scarcity issue, pan-specific methods have been proposed and have achieved significant improvement in terms of prediction performance [26]. In these models, binding peptides of different alleles are all combined to train a single prediction model for all HLA alleles. Typically, a pan-specific model uses binding affinity data from multiple alleles for training and could predict peptide binding affinity for the alleles that may have or have not appeared in the training data. The key idea behind pan-specific models is that besides encoding the peptide in a proper way for the prediction model, the peptide-HLA binding context/environment is also represented so that the machine learning models could be trained on all available peptide-HLA binding samples [26]. In other words, both the peptide and the HLA protein are encoded as input to the pan-specific models to train the prediction models. So far, a number of pan-specific models have been proposed for both HLA class I and class II alleles [26]. Among them, NetMHCPan, PickPocket and Kim et al’s work are recently proposed pan-specific HLA-peptide binding prediction models trained on the large amount of HLA class I binding affinity data.

NetMHCPan is the first pan-specific binding affinity prediction algorithm that takes a large number of peptide-HLA binding samples of different HLA alleles for model training and obtained state-of-the-art performance [6]. NetMHCPan proposed a novel pseudo sequence encoding method to represent the binding context, in which an HLA sequence is reduced to a pseudo amino acid sequence of length 34. Each amino acid in this pseudo sequence is selected if it is in contact with the peptide within 4.0 Å (as shown in Figure 1). The interaction map in Figure 1 is extracted based on a representative set of HLA structures with nonamer peptides. Actually, this extracted 34-length pseudo sequence is a list of location indexes of amino acids in the HLA sequence. For a given HLA sequence, only 34 corresponding residues are encoded as input. In NetMHCPan, a HLA-peptide binding sample is represented as a 43-length residue sequence (9 from the peptide and 34 from the HLA). This sequence is then encoded in three different ways: one-hot encoding, BLOSUM50, and a mixture of both. The encoded input is then used to train multiple feedforward neural networks with 22 to 86 hidden neurons. Then the network with the highest prediction performance (lowest square error) on the test set was selected as the final prediction model [6]. This pseudo sequence encoding approach for pan-specific modeling has also been used in PickPocket [25] and Kim’s algorithm [4], but with different machine learning algorithms for model training. In PickPocket, position-specific scoring matrices (PSSMs) are first derived from peptides data. Then extract the position-specific vectors from the PSSMs in association with pseudo-sequence to construct a pocket library. Each pocket library entry is characterized by nine pairs, where each pair consists of a list of pocket amino acids and a specificity vector.

**Figure 1.**
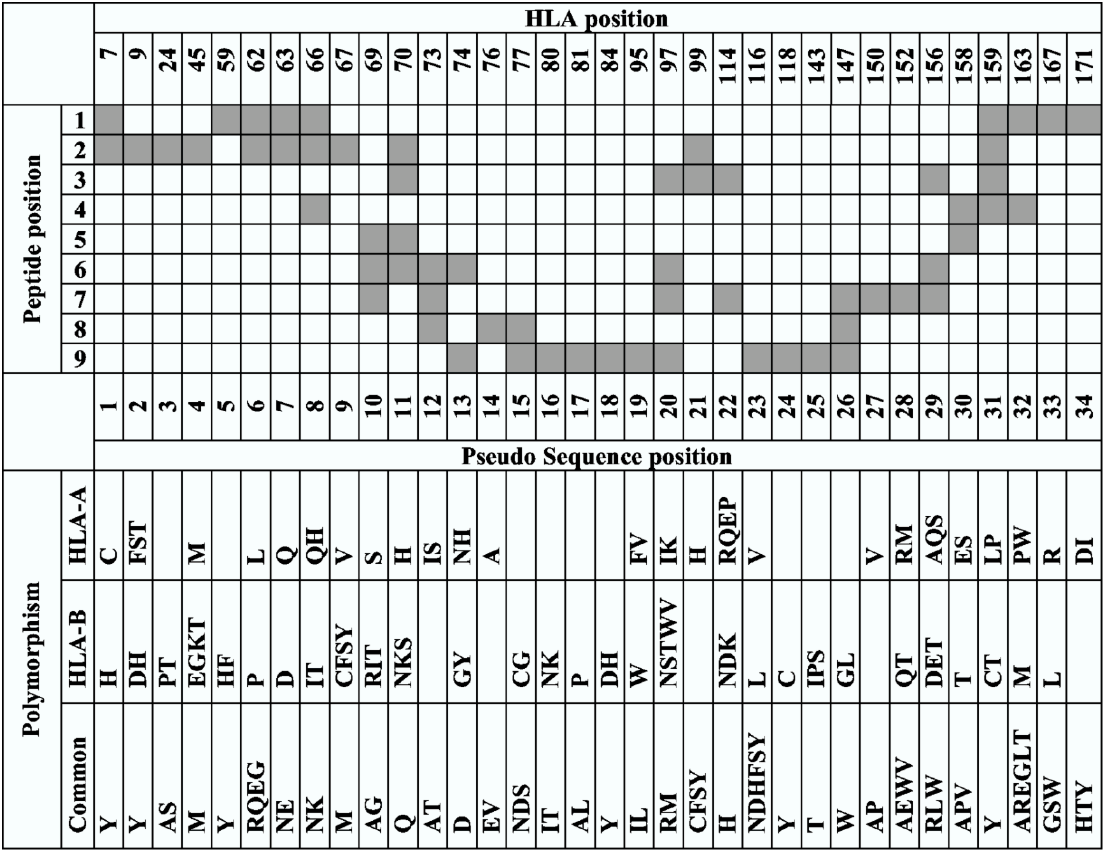
Interaction map of the HLA pseudo sequence in NetMHCPan. Reproduced from original paper.

Deep convolutional neural networks (DCNN) are powerful deep learning models and have been successfully applied in many bioinformatics problems such as DNA-binding prediction and CpG island binding prediction [1, 2]. This technique has also been applied to peptide-MHC binding prediction [4,7,23]. Among them, Kim et al. proposed a pan-specific DCNN model for peptide-MHC class I binding prediction [4], in which the peptide-binding context is encoded using NetMHCPan’s contact residue sequence method and the DCNN model is trained as a 26-layer classifier.

To our best knowledge, the pseudo sequence encoding proposed in NetMHCPan is currently the only binding context encoding method in pan-specific peptide-HLA class I binding prediction, which has achieved state-of-the-art performance in the public benchmark study [22]. However, this encoding method has its potential limitations: 1) its interaction map extraction step relies on available MHC-peptide bound complex structures, which may not be always available, especially considering the high polymorphism of HLA proteins; 2) the 34 contact residues of the encoding is empirical and only covers part of the whole HLA sequence.

In this paper, we propose DeepSeqPan (Figure 2), an end-to-end deep learning model trained on pairs of one-hot encoded raw peptides and HLA sequences, which make it possible to train pan-specific HLA-peptide binding prediction model without the three dimensional structural data. Evaluation on the independent IEDB benchmark datasets showed that our proposed model achieved highly competitive performance on many HLA alleles. Our contributions can be summarized as follows:

1) we proposed a novel DCNN architecture for pan-specific HLA binding prediction, in which the peptide-HLA binding context is learned by the network itself. 2) Our DeepSeqPan does not rely on any structure information 3) Designed as a multi-task learning problem, our DCNN predicts both IC_50_ values and binding probabilities at the same time. 4) Our pan-specific prediction model has demonstrated high generalization capability across alleles. In a blind testing, a model was trained only on HLA-A and -B alleles and achieved very good performance when tested on HLA-C alleles.

**Figure 2.**
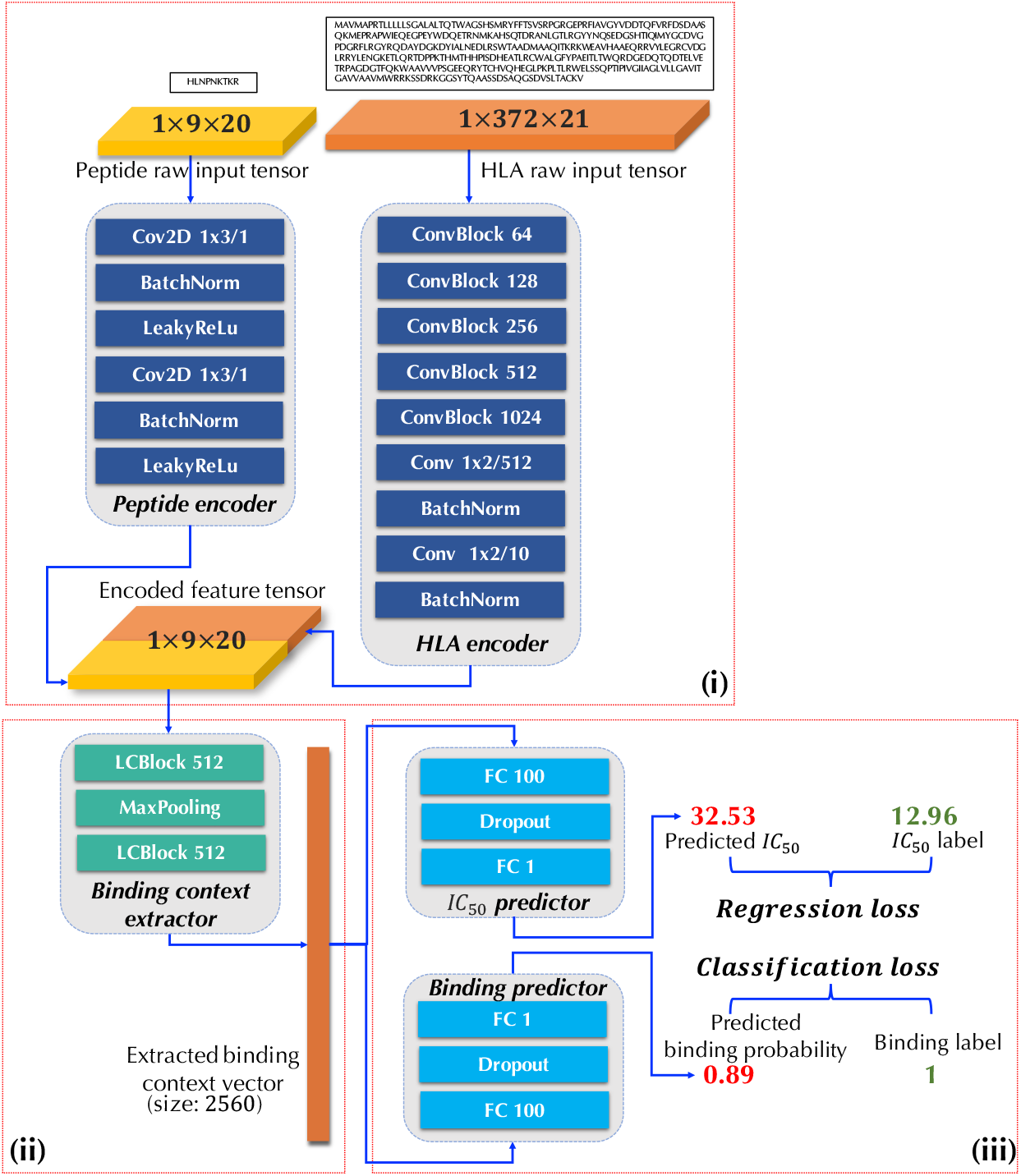
DeepSeqPan Network Structure, (i) Peptide and HLA encoders, (ii) Binding context extractor, (iii) Affinity and binding predictors.

## Method

### Dataset

The training dataset BD2013 is downloaded from the widely used IEDB [24] database ^2^. All training samples are labeled with IC_50_ binding affinity values. The testing dataset is downloaded from IEDB’s weekly benchmark dataset ranging from 2014-03-21 to 2018-01-26 ^3^. To address the concern that duplicate peptides may exist in both the training and testing data downloaded from IEDB, we removed all duplicate peptides from the testing dataset. The aligned HLA sequences were obtained from IMGT/HLA database [19]. We trained our model on 9-length peptides binding to HLA-A, HLA-B and HLA-C alleles with available HLA sequences. Totally, the training dataset contains 121,787 peptide-HLA binding peptides covering 42 HLA-A alleles (72618 samples), 49 HLA-B alleles (46915 samples) and 10 HLA-C alleles (2254 samples). The detailed information of the training and testing data are listed in Supplementary File.

### Sequence Encoding

In our DeepSeqPan model, each input is a HLA-peptide pair. For both the peptide and the HLA in an input, we use the naive one-hot encoding according to amino acids’ locations in sequences. A 9-length peptide sequence is encoded into a 2D tensor with dimension 1 (height) × 9 (width) × 20 (channel) where the last dimension is the number of channels and each channel represents one of 20 amino acids. Figure 3 illustrates the encoded peptide HLNPNKTKR as a 2D tensor with dimension 1 × 9 × 20. Since HLA sequences have variable lengths, we chose the maximum length 372 as the fixed dimension. Then we encode each aligned HLA sequence into a 2D tensor with dimension 1 × 372 × 21. The extra channel represents gaps in HLA sequences shorter than 372.

**Figure 3.**
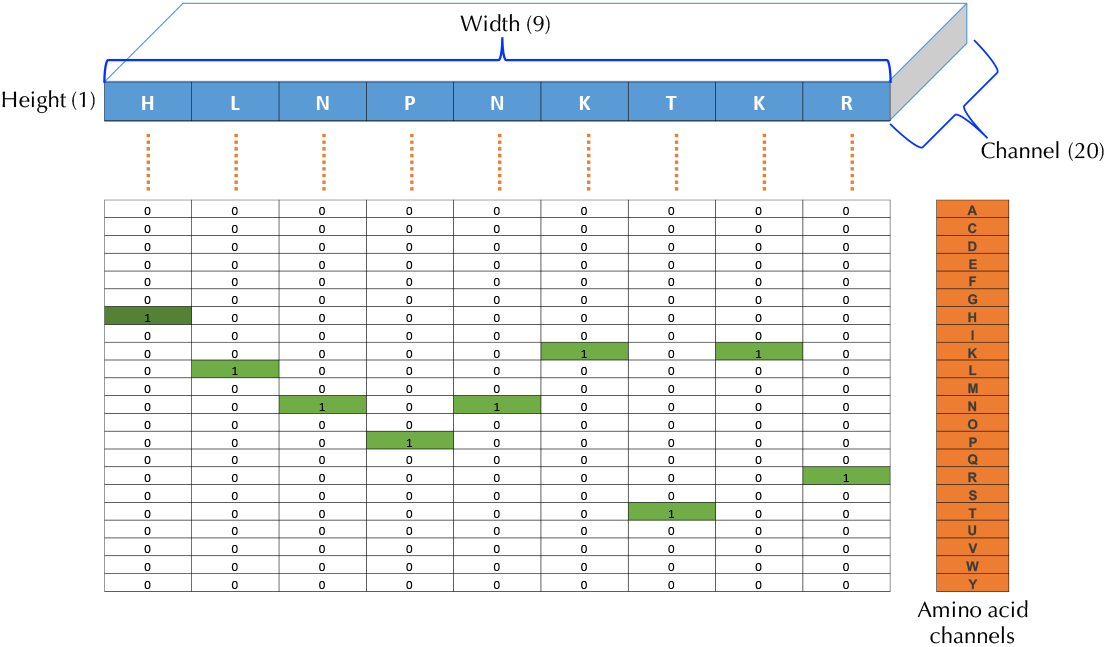
Peptide encoding example. Sequence HLNPNKTKR is encoded into a 2D tensor with dimension 1 (*height*) × 9(*width*) × 20(*channel*). Each of 20 channels represents one amino acid type and we set a channel value to 1 if the corresponding amino acid appears at this location of the input sequence.

### The deep neural network model of DeepSeqPan

#### Architecture

As shown in Figure 2, the DeepSeqPan network consists of three parts:

i. **Peptide and HLA encoders.** The peptide and HLA encoders convert a pair of one-hot encoded peptide and HLA sequences into two tensors with a unified dimension 1 × 9 × 10. The output tensors of two encoders are concatenated along the channel axis to generate an encoded feature tensor with dimension 1 × 9 × 20. Then this concatenated tensor will be fed into the binding context extractor in (ii). The purpose of these two encoders is extracting high-level features from raw sequences and encoding them into a feature tensor. Different from the 34 pseudo amino acid sequence encoding approach in [14], the features and information stored inside this feature tensor are learned by the deep neural network automatically with its end-to-end training framework. The encoder of the peptide consists of two blocks of convolutional, batch normalization and LeakyReLu layers. As for the HLA encoder whose input sequence is much longer than the peptide, we used a network configuration similar to the VGG network [20].
ii. **Binding context extractor.** The extractor takes into the encoded feature tensor from (i) and outputs a 2560-dimension vector. This vector is actually the binding context between a peptide and a HLA. This binding context extractor will be optimized automatically in the training stage through the backpropagation algorithm and the extraction of the binding context is done by the network itself without human involvement. Especially, in this extractor we use Locally Connected layers (as illustrated *LCBlock* in Figure 2) instead of standard convolutional layers with weights sharing. The reason is that the encoded high-level features in the feature tensor is position related, i.e. in the encoded feature tensor with length 9, an extracted feature 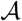 located at position 1 should have different effect as it appears at position 7. Locally connected layers has the capability to capture features at specific locations since its filters at different locations do not share weights, which has been proved to be powerful in DeepFace [21].
iii. **Affinity and binding predictors.** Another novel design of DeepSeqPan is that at the output layer, both the binding probabilities and the IC_50_ value are used as output in final stage (iii). This is different from all other DCNN based MHC binding prediction algorithms [4, 7, 23] which outputs either the binding probabilities or IC_50_ values. This design is not a captain’s call. Actually, at first when we were training the DeepSeqPan that only predicts IC_50_ values, we found it was very hard to train the network with very slow convergence. So we added the binding probability predictor as an additional source of supervision signal with the expectation that the backpropagation algorithm can train the network easier by taking advantage of two types of losses: the classification loss and regression loss. Note that we can calculate the binary binding probability for the training samples from their IC_50_ binding affinity values via Equation (4). The underlying relationship between regression outputs and classification outputs is built up naturally. In the training stage, the network needs to learn this underlying relationship in order to reduce the total loss. In that case, we argue that the classification predictor plays as a regularizer by forcing the network to predict a more accurate IC_50_ values.

We went through a grid search over hyperparameters setting and the detailed configurations of the layers/blocks and associated hyperparameters are described in the Supplementary File.

#### Loss Function

The overall loss 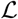 is the sum of the regression loss 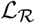 and the classification loss 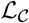 as illustrated in Equation (1).

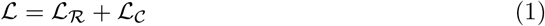

For IC_50_ predictor, we use mean squared error (MSE, Equation (2)) as the loss function and for the binary binding predictor, the binary cross entropy loss (Equation (3)) is used.

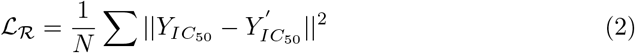

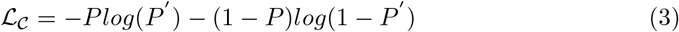

To get binary binding labels, we use standard 500 nM threshold to convert a IC_50_ value label into a binding label:

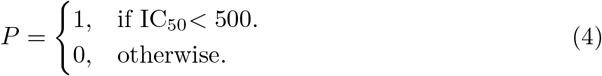

#### Training

We randomly split all training samples into a training set and a validation set following 4:1 ratio. Stochastic gradient descent (SGD) is employed as the optimization algorithm enabled with momentum and learning rate decay. The initial learning rate is 0.001 and the momentum factor is 0.8. It is scheduled to halve the learning rate if validation loss hasn’t improved within 5 epochs. The minimum learning rate is set to 0.00001. The training process stops if the validation loss has not improved within 20 epochs. We used Keras [3] deep learning framework to implement our DeepSeqPan algorithm.

### Metrics and label preprocessing

Area under the curve (AUC) and Spearman’s rank correlation coefficient (SRCC) are used as evaluation metrics to compare with the public benchmark results at IEDB website [22].

In pan-specific binding prediction modeling, the IC_50_ values of the peptides span a large range [0, 80000]. To avoid gradient explosion issue in neural network training, we convert IC_50_ to logIC_50_ via Equation 5. The logIC_50_ are then used as labels during training. During inference stage, we convert the prediction results back to IC_50_ values.

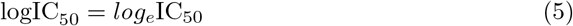

## Results and discussion

### Cross-validation on the training dataset

Standard five-fold cross-validation experiments were applied to the different alleles and their combinations in the training dataset. The performance is then measured in area under a curve (AUC) and Spearman’s rank correlation coefficient (SRCC). Since our network outputs both IC_50_ affinity values and binding probabilities, we evaluated the performances on both outputs separately, in terms of classification performance and regression performance.

Table 1 shows the 5-fold cross-validation results of our algorithm on the training dataset. When all the samples are included, DeepSeqPan achieved a high AUC of 0.94 for regression on (IC_50_) and an AUC of 0.94 for binary binding classification. The corresponding SRCC is 0.73 (IC_50_) and 0.70 (binary binding) respectively. When evaluated over the samples of HLA-A, -B and -C alleles separately, all the AUC scores are above 0.90 and SRCC scores are all above 0.70. More comprehensive allele-specific evaluation results are reported in Supplementary Files. The cross-validation results indicate that DeepSeqPan can be trained to achieve high-performance HLA binding peptide prediction with strong generalization capability.

**Table 1.** Five-fold cross validation results

| Alleles | Sequence Count | $IC_{50}$ | | Binary binding | |
| --- | --- | --- | --- | --- | --- |
|  |  | AUC | SRCC | AUC | SRCC |
| All alleles | 121787 | 0.94 | 0.73 | 0.94 | 0.70 |
| HLA-A alleles | 72618 | 0.94 | 0.75 | 0.94 | 0.73 |
| HLA-B alleles | 46915 | 0.94 | 0.68 | 0.94 | 0.64 |
| HLA-C alleles | 2254 | 0.89 | 0.70 | 0.89 | 0.69 |

### Evaluation on benchmark dataset

To evaluate how our DeepSeqPan performs compared to other HLA-peptide binding prediction algorithms, we applied it to the public IEDB weekly benchmark dataset upon which a set of top algorithms have been evaluated with published results.

We trained a single DeepSeqPan model on all 9-length peptides in the training dataset that bind to HLA-A, -B and -C alleles. Then this trained model was evaluated on all available IEDB weekly benchmark dataset [22]. As we inform before, the IEDB benchmark dataset has been filtered by removing duplicate samples. We compared the performance of DeepSeqPan with those of pan-specific models: NetMHCPan(2.8) [6] and PickPocket [25], the performances of allele-specific models: SMM [17], NetMHC(3.4) [12], ARB [22], MHCflurry [16] and AMMPMBEC [22], and those of ensemble models (results are based on several different models): IEDB Consensus [22] and NetMHCcons [9]. All the prediction results of compared models in Table 2 are provided in the original benchmark dataset downloaded from the IEDB benchmark website.

**Table 2.**
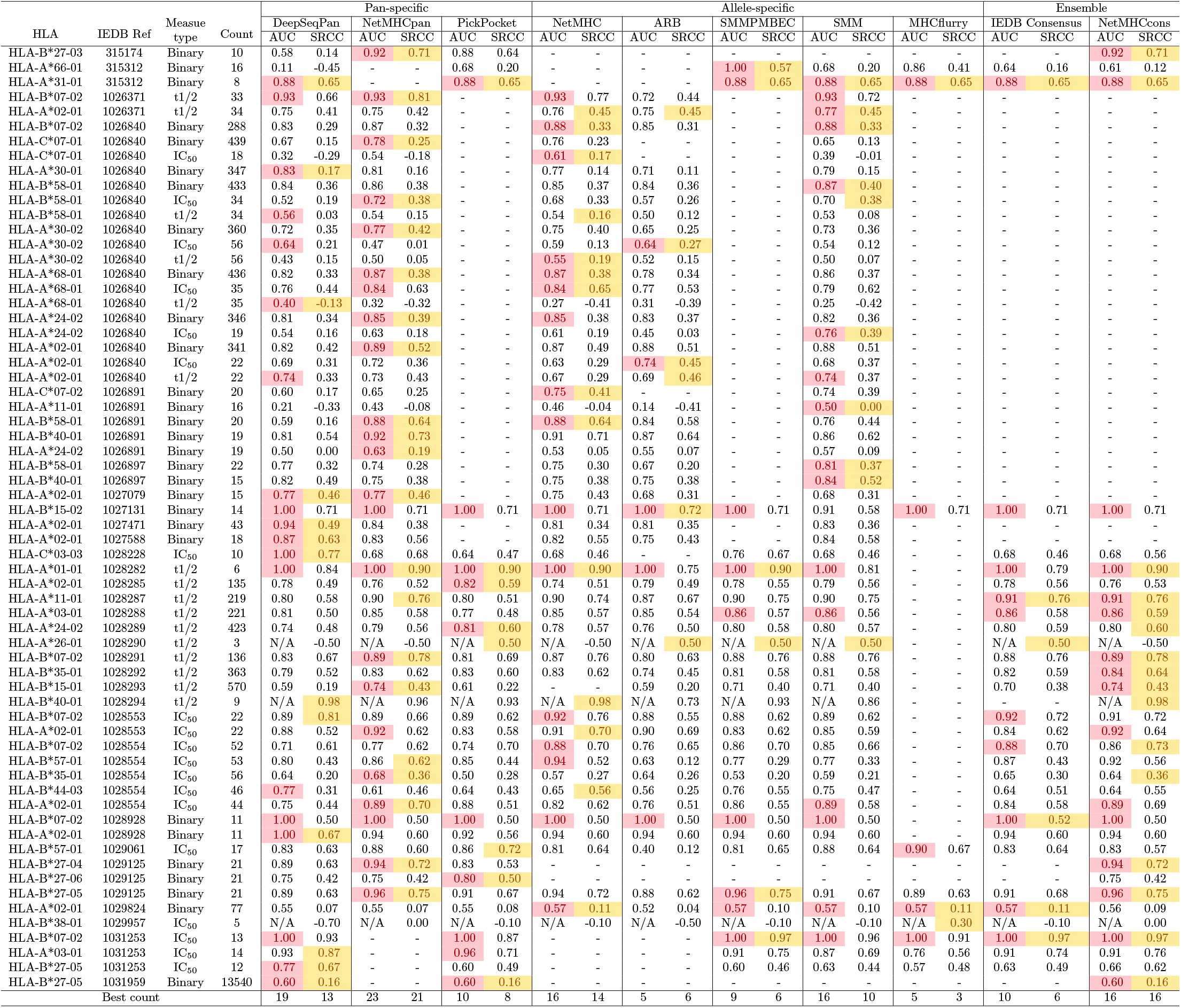
Evaluation on benchmark database (Red/Yellow highlighted number(s) are best AUC/SRCC scores in that record). Sorted by IEDB Ref ID.

Table 2 summarized the performance of different algorithms on 64 testing datasets from IEDB benchmark database. For each dataset, we highlighted the highest AUC scores in yellow and highest SRCC scores in pink and then counted the number of datasets upon which each algorithm achieved the highest scores and put them at the last row of the table.

First we compared the performance of DeepSeqPan with other pan-specific models and ensemble model based on pan-specific models. We found that DeepSeqPan showed highly competitive performance by achieving the highest AUC scores in 19 datasets out of total 64 testing datasets. This is only second to NetMHCPan, which outperforms others in 23 datasets and uses structural information of HLA-peptide complex structure information. However, in 45 datasets that DeepSeqPan didn’t achieve the highest AUC scores, there are 28 datasets on which the AUC scores of DeepSeqPan are very close to the highest AUC scores within a small margin around 0.1. Our algorithm also outperforms another pan-specific model, PickPocket, which has the highest AUC scores over 10 datasets. Our method also beats the best ensemble method NetMHCcons with the best performance on 16 datasets, which is based on fusion of NetMHC, NetMHCPan and PickPocket [9]. In terms of SRCC, DeepSeqPan obtained the highest scores on 13 datasets compared to 21 of NetMHCPan, 16 of NetMHCcons, and 14 of NetMHC.

Overall, the benchmark evaluation results showed that DeepSeqPan is highly competitive in HLA-peptide binding prediction with 2nd most highest AUC scores and 4th highest SRCC scores. This is impressive considering that our model was trained only on raw amino acid sequence without relying on any structure or amino acid properties information. From Table 2, it can be found that different pan-specific and allele-specific methods have the best performance on datasets of various alleles, which implies the good performance of the ensemble methods such as NetMHCcons since they make prediction via combining results from multiple methods. Our proposed DeepSeqPan could thus be a complementary tool for existing pan-specific models and it is promising to include it into the state-of-the-art ensemble prediction models to improved their performance.

### Comparison with other DCNN models

To the best of our knowledge, Kim el at.’s work [4] is the only pan-specific model that employs DCNN architecture beside our proposed DeepSeqPan. It uses NetMHCPan’s pseudo sequence encoding for binding context modeling, in which a pair of peptide-HLA binding sample is encoded into a 9 (height) × 34 (width) × 18 (channel) 2D tensor. Each “pixel” in this 2D tensor represents a contacting pair of a peptide reside and a HLA residue. For two contacting residues, 9 physicochemical properties are used for each one and in total 18 values are encoded in channels. Their network structure is VGG-like and consists of 26 layers. They trained their model with binding samples on HLA-A and HLA-B alleles and it used the same dataset BD2013 as we did. To compare the performance of our DeepSeqPan with Kim’s method, we evaluated the benchmark dataset with their online server4 on all its supporting alleles (HLA-A and HLA-B alleles). In total, we evaluated 54 benchmark dataset on Kim’s server and compared with ours obtained in previous benchmark evaluation and the binary prediction outputs were used to compare. Since Kim’s model was trained as a classifier, we calculated AUC scores for each testing dataset and in Table 3 we showed the average AUC scores measured based on all HLA-A or HLA-B testing dataset respectively (Detailed performance on each dataset is listed in Supplementary Files). Out of all 54 benchmark dataset, Kim’s model and our model both got an average AUC of 0.76. For HLA-A datasets, two model also obtained same average AUC of 0.74. Our model slightly out performed Kim’s model on HLA-B alleles with an average AUC of 0.80. Overall, two models achieved similar performance and in terms of performance on each allele as shown in Supplementary Files, two models obtained better performance on different sets of HLA alleles and none can dominate the other model.

**Table 3.** Comparison of Kim’s model and DeepSeqPan

| HLA | # of Records | Avg. AUC |  |
| --- | --- | --- | --- |
|  |  | Kim’s model | DeepSeqPan (Binary) |
| All | 54 | 0.76 | 0.76 |
| HLA-A | 30 | 0.74 | 0.74 |
| HLA-B | 24 | 0.79 | 0.80 |

### Generalization of DeepSeqPan to binding predictions of new HLA alleles

One major advantage of pan-specific models over allele-specific models is that it can give prediction on HLA alleles that are not included in the training dataset. This is especially useful for HLA alleles without any samples with known binding affinity values. In order to evaluate this extrapolation capability of DeepSeqPan, we setup an experiment in which the models are trained on samples that only bind to HLA-A and HLA-B alleles. We call this trained model *Blind model* since it does not include any HLA-C samples in its training data (so it is blind to the test samples that binds HLA-C). Then we predicted the binding of the peptide samples that bind to HLA-C alleles with this trained Blind model. All samples are taken from the training dataset (i.e. BD2013) such that we can compare this model’s performance with that of 5-fold cross-validation.

The comparison results are shown in Table 4. The columns under *Blind* are performance results of the *Blind model.* Columns under *Cross Validation* are results obtained from previous 5-fold cross-validation. We reported two methods’ performance metrics on all HLA-C samples and on samples of each HLA-C allele. The Blind model achieved an AUC of 0.64/0.66 and SRCC of 0.26/0.29 as measured in IC_50_/Binary on all 2254 HLA-C samples. Over individual HLA-C alleles, the Blind model achieved AUC scores of 0.5-and-above over 6 out of 10 HLA-C alleles when measured in both IC_50_ and binary. This demonstrated that our DeepSeqPan has the capability to generalize across HLA alleles. When compared with cross-validation results, as expected, the performance of the Blind model dropped around 0.1 ~ 0.4 in terms of the AUC scores. However, on allele HLA-C*12:03, the Blind model performed better than the cross-validation model with higher AUC scores, which is beyond expectation. Considering the blind training process of DeepSeqPan here in which the HLA-C samples are not included in the training samples, the overall performance of the Blind model is impressive and it shows the HLA encoder in DeepSeqPan could capture the general high-level features of HLA alleles even when trained only with samples of HLA-A and -B alleles. The results showed that our proposed neural network model has good generalization capability across alleles.

**Table 4.** Performance comparison on affinity prediction of samples binding to HLA-C alleles

| HLA | Count | Blind |  |  |  | Cross Validation |  |  |  |
| --- | --- | --- | --- | --- | --- | --- | --- | --- | --- |
|  |  | IC<sub>50</sub> |  | Binary |  | IC<sub>50</sub> |  | Binary |  |
|  |  | AUC | SRCC | AUC | SRCC | AUC | SRCC | AUC | SRCC |
| All C alleles | 2254 | 0.64 | 0.26 | 0.66 | 0.29 | 0.89 | 0.70 | 0.89 | 0.69 |
| HLA-C*03:03 | 154 | 0.60 | 0.27 | 0.59 | 0.27 | 0.80 | 0.59 | 0.79 | 0.58 |
| HLA-C*04:01 | 520 | 0.36 | 0.07 | 0.32 | 0.00 | 0.52 | -0.09 | 0.58 | -0.15 |
| HLA-C*05:01 | 172 | 0.46 | 0.00 | 0.47 | 0.00 | 0.91 | 0.74 | 0.92 | 0.75 |
| HLA-C*06:02 | 301 | 0.58 | 0.30 | 0.60 | 0.32 | 0.89 | 0.74 | 0.89 | 0.74 |
| HLA-C*07:01 | 220 | 0.49 | -0.02 | 0.49 | 0.00 | 0.84 | 0.61 | 0.84 | 0.61 |
| HLA-C*07:02 | 140 | 0.63 | 0.15 | 0.64 | 0.17 | 0.74 | 0.42 | 0.73 | 0.41 |
| HLA-C*08:02 | 87 | 0.48 | -0.10 | 0.46 | -0.13 | 0.64 | 0.28 | 0.63 | 0.25 |
| HLA-C*12:03 | 172 | 0.65 | 0.18 | 0.65 | 0.18 | 0.62 | 0.28 | 0.61 | 0.29 |
| HLA-C*14:02 | 235 | 0.54 | 0.09 | 0.57 | 0.11 | 0.67 | 0.28 | 0.67 | 0.29 |
| HLA-C*15:02 | 253 | 0.61 | 0.14 | 0.63 | 0.17 | 0.79 | 0.52 | 0.80 | 0.55 |

### The binding context vector: consistency and capability

One of key design features of our DeepSeqPan model (Figure 2) is the binding context vector aiming to capture high-level features that determine whether a peptide and a HLA will bind or not and if so, how strong the binding is. Another key feature of our model is the dual outputs of the model: the binding affinity output and the binary binding probability output.

Since the binding context vector is used as input for both predicted outputs, it should be consistent for both the IC_50_ predictor and the binary binding predictor in (iii): for the same binding context vector, both predictors should give consistent outputs. In other words, higher binding probability should correspond to higher binding affinity values. To verify this consistency, we inspected IC_50_ and binary prediction outputs of all samples from previous cross-validation and benchmark evaluation experiments. The analysis results are shown in Table 5. First row of the table lists the numbers of evaluated samples in the cross-validation and benchmark evaluation experiments with 121, 787 in cross-validation and 19, 741 in benchmark evaluation. The second row shows the number of consistent outputs. Given a sample pair of a peptide and an HLA, we mark its predicted IC_50_ value and predicted binding probability consistent if both values indicate binding or not binding. An IC_50_ value of <500 nM indicates binding and a binding probability of 0.5 or greater means the binding state. From the table we can observe that high consistency exists between the regression and classification outputs. For cross-validation experiments, the percentage of consistent outputs is 95.81% and for benchmark evaluation experiments, this percentage is 86.14%.

**Table 5.** Consistency inspection results

|  | Cross Validation | Benchmark Evaluation |
| --- | --- | --- |
| Total samples | 121787 | 19741 |
| Consistent pred. | 116688 (95.81%) | 17004 (86.14%) |
| Correct IC<sub>50</sub> pred. | 108064 (88.73%) | 11690 (59.21%) |
| Correct Binary pred. | 107239 (88.05%) | 10487 (53.12%) |

In last 2 rows of Table 5, we reported the number of correct predictions measured with the IC_50_ predictions and the binary prediction respectively. A predicted IC_50_ value or a predicted binary binding probability value is marked as a correct prediction if its real label and the prediction value indicate the same binding state: binding or not binding. Given a sample, it will be marked as binding if its IC_50_ value is greater than the threshold (500 nM). If the sample binding affinity is labeled with t1/2 type, an affinity value of < 120 indicates the binding state. For binary binding labels, a binary label of 1 means it is binding while a value of 0 means no binding. For the predicted binary binding probability, a probability value of > 0.5 means binding. From Table 5, we found that both affinity and binary binding outputs obtained accuracies greater than 88% in cross-validation experiments. In benchmark experiments, the accuracy rate is 59% for IC_50_ predictions and binary predictions have an accuracy of 53%. The results showed that the consistency between IC_50_ predictions and binary predictions is high, which means that the binding context vector extracted by DeepSeqPan contains common effective features for determining binding states.

Table 1, 2 and 5 together showed that the binding context vector learned via the end-to-end learning framework of the DCNN is predictor-independent and has captured information related to HLA-peptide binding. Its effectiveness in HLA binding prediction may be explained by its capability to capture position related information such as when there’s an animo acid A in HLA at position 37 and at the same time there’s an amino acid L in peptide at position 5, the binding affinity is high. This binding feature extraction is similar to DCNN architectures like VGG16 [20], ResNet [5] of computer vision in which an input image will be represented as a high-dimension vector (4096 in VGG16 and 1000 in ResNet) in the final stage of the neural network. Due to time and hardware resource constraints, our current version of DeepSeqPan only uses one-hot encoding information. No other information such as physical properties of amino acids are encoded into input tensors, which however can further improve the performance of DeepSeqPan if properly encoded by capturing more relevant and rich binding contexts.

## Conclusion

In this work, we designed, DeepSeqPan, a novel deep convolutional neural network model for pan-specific HLA-peptide binding affinity prediction. This model is characterized by its capability of binding prediction with only the raw amino acid sequences of the peptide and the HLA, which makes it applicable to HLA-peptide binding prediction for HLA alleles without structural information. This is achieved by a novel sequence based encoding of the peptide-HLA binding context, a binding context feature extractor, and the dual outputs with both binding affinity and binding probability predictions. Extensive evaluation of DeepSeqPan on public benchmark experiments showed that our model achieves state-of-the-art performance on an variety of HLA allele datasets.

Our model contributes to the study of MHC-peptide binding prediction in a few special ways. First, our experiments showed that it is possible to extrapolate the binding prediction capability to unseen HLA alleles, which is important for pan-specific models. Second, our sequence-only based binding context encoding is complementary to to the pseudo sequence encoding, which is currently the only encoding method used in pan-specific models for class I MHC-peptide binding affinity prediction. This has the potential to further improve the state-of-the-art prediction models such as the pan-specific model NetMHCSpan. It showed the importance of sufficient amount of training data to achieve high prediction performance for deep learning models.

Our current work can be further improved in a number of ways. First, in this work, only one-hot encoding is used for representing the input peptide and HLA protein. However, this can be improved by properly encoding more features such as physicochemical properties of amino acids into input tensors. Moreover, our proposed sequence-based DCNN architecture for protein-peptide binding is universal and can be adapted to other similar binding problems such as protein-DNA, protein-RNA and protein-ligand/drug bindings.

## Acknowledgments

We gratefully acknowledge the support of NVIDIA Corporation with the donation of the Titan X GPU used for this research.

## Author Contribution Statement

Z.L and J.H wrote the main manuscript text. Y.C, A.Z and Z.X prepared all data tables. A.N prepared all figures. All authors reviewed the manuscript.

## Data availability

The implementation code and trained models are freely available at https://github.com/pcpLiu/DeepSeqPan

## Footnotes

1 https://www.ebi.ac.uk/ipd/imgt/hla/stats.html

2 http://tools.iθdb.org/main/datasθts/

3 http://tools.iθdb.org/auto_bench/mhci/weekly/

4 http://jumong.kaist.ac.kr:8080/convmhc

## References

1. B. Alipanahi, A. Delong, M. T. Weirauch, and B. J. Frey. Predicting the sequence specificities of dna-and rna-binding proteins by deep learning. Nature biotechnology, 33(8):831, 2015.

2. C. Angermueller, H. J. Lee, W. Reik, and O. Stegle. Deepcpg: accurate prediction of single-cell dna methylation states using deep learning. Genome biology, 18(1):67, 2017.

3. F. Chollet et al. Keras. https://keras.io, 2015.

4. Y. Han and D. Kim. Deep convolutional neural networks for pan-specific peptide-mhc class i binding prediction. BMC bioinformatics, 18(1):585, 2017.

5. K. He, X. Zhang, S. Ren, and J. Sun. Deep residual learning for image recognition. In Proceedings of the IEEE conference on computer vision and pattern recognition, pages 770–778, 2016.

6. I. Hoof, B. Peters, J. Sidney, L. E. Pedersen, A. Sette, O. Lund, S. Buus, and M. Nielsen. Netmhcpan, a method for mhc class i binding prediction beyond humans. Immunogenetics, 61(1):1, 2009.

7. J. Hu and Z. Liu. Deepmhc: Deep convolutional neural networks for high-performance peptide-mhc binding affinity prediction. bioRxiv, page 239236, 2017.

8. L. Jacob and J.-P. Vert. Efficient peptide-mhc-i binding prediction for alleles with few known binders. Bioinformatics, 24(3):358–366, 2007.

9. E. Karosiene, C. Lundegaard, O. Lund, and M. Nielsen. Netmhccons: a consensus method for the major histocompatibility complex class i predictions. Immunogenetics, 64(3):177–186, 2012.

10. S. Lata, M. Bhasin, and G. P. Raghava. Mhcbn 4.0: A database of mhc/tap binding peptides and t-cell epitopes. BMC research notes, 2(1):61, 2009.

11. G. Liu, D. Li, Z. Li, S. Qiu, W. Li, C.-c. Chao, N. Yang, H. Li, Z. Cheng, X. Song, et al. Pssmhcpan: a novel pssm-based software for predicting class i peptide-hla binding affinity. Giga Science, 6(5):1–11, 2017.

12. C. Lundegaard, K. Lamberth, M. Harndahl, S. Buus, O. Lund, and M. Nielsen. Netmhc-3.0: accurate web accessible predictions of human, mouse and monkey mhc class i affinities for peptides of length 8–11. Nucleic acids research, 36(suppl_2):W509–W512, 2008.

13. H. Luo, H. Ye, H. W. Ng, S. Sakkiah, D. L. Mendrick, and H. Hong. snebula, a network-based algorithm to predict binding between human leukocyte antigens and peptides. Scientific reports, 6:32115, 2016.

14. M. Nielsen and M. Andreatta. Netmhcpan-3.0; improved prediction of binding to mhc class i molecules integrating information from multiple receptor and peptide length datasets. Genome medicine, 8(1):33, 2016.

15. M. Nielsen, C. Lundegaard, T. Blicher, B. Peters, A. Sette, S. Justesen, S. Buus, and O. Lund. Quantitative predictions of peptide binding to any hla-dr molecule of known sequence: Netmhciipan. PLoS computational biology, 4(7):e1000107, 2008.

16. T. O’Donnell, A. Rubinsteyn, M. Bonsack, A. Riemer, and J. Hammerbacher. Mhcflurry: open-source class i mhc binding affinity prediction. bioRxiv, page 174243, 2017.

17. B. Peters and A. Sette. Generating quantitative models describing the sequence specificity of biological processes with the stabilized matrix method. BMC bioinformatics, 6(1):132, 2005.

18. H.-G. Rammensee, J. Bachmann, N. P. N. Emmerich, O. A. Bachor, and S. Stevanović. Syfpeithi: database for mhc ligands and peptide motifs. Immunogenetics, 50(3–4):213–219, 1999.

19. J. Robinson, J. A. Halliwell, J. D. Hayhurst, P. Flicek, P. Parham, and S. G. Marsh. The ipd and imgt/hla database: allele variant databases. Nucleic acids research, 43(D1):D423–D431, 2014.

20. K. Simonyan and A. Zisserman. Very deep convolutional networks for large-scale image recognition. arXiv preprint arXiv:1409.1556, 2014.

21. Y. Taigman, M. Yang, M. Ranzato, and L. Wolf. Deepface: Closing the gap to human-level performance in face verification. In Proceedings of the IEEE conference on computer vision and pattern recognition, pages 1701–1708, 2014.

22. T. Trolle, I. G. Metushi, J. A. Greenbaum, Y. Kim, J. Sidney, O. Lund, A. Sette, B. Peters, and M. Nielsen. Automated benchmarking of peptide-mhc class i binding predictions. Bioinformatics, 31(13):2174–2181, 2015.

23. Y. S. Vang and X. Xie. Hla class i binding prediction via convolutional neural networks. Bioinformatics, 33(17):2658–2665, 2017.

24. R. Vita, J. A. Overton, J. A. Greenbaum, J. Ponomarenko, J. D. Clark, J. R. Cantrell, D. K. Wheeler, J. L. Gabbard, D. Hix, A. Sette, et al. The immune epitope database (iedb) 3.0. Nucleic acids research, 43(D1):D405–D412, 2014.

25. H. Zhang, O. Lund, and M. Nielsen. The pickpocket method for predicting binding specificities for receptors based on receptor pocket similarities: application to mhc-peptide binding. Bioinformatics, 25(10):1293–1299, 2009.

26. L. Zhang, K. Udaka, H. Mamitsuka, and S. Zhu. Toward more accurate pan-specific mhc-peptide binding prediction: a review of current methods and tools. Briefings in bioinformatics, 13(3):350–364, 2011.

